# Antibody interference by a non-neutralizing antibody abrogates humoral protection against *Plasmodium* liver stage

**DOI:** 10.1101/2020.09.15.298471

**Authors:** Kamalakannan Vijayan, Ramyavardhanee Chandrasekaran, Olesya Trakhimets, Samantha L. Brown, Nicholas Dambrauskas, Meghan Zuck, Ganesh Ram R. Visweswaran, Alexander Watson, Andrew Raappana, Sara Carbonetti, Laurel Kelnhofer-Millevolte, Elizabeth K.K. Glennon, Rachel Postiglione, D. Noah Sather, Alexis Kaushansky

## Abstract

Both subunit and attenuated whole sporozoite vaccination strategies against *Plasmodium* infection have shown promising initial results in malaria-naïve westerners but exhibited less efficacy in malaria-exposed individuals in endemic areas. It has been hypothesized that preexisting immunity to malaria represents a significant roadblock to the development of a protective vaccine. Here, we demonstrate proof-of-concept that non-neutralizing antibodies (nNAb) can directly interfere with protective anti-PyCSP humoral responses. We developed and characterized a novel monoclonal antibody, RAM1, against the *P. yoelii* sporozoite major surface antigen, circumsporozoite protein (CSP). Unlike the canonical *Py*CSP repeat domain binding and neutralizing antibody (NAb) 2F6, RAM1 does not inhibit sporozoite traversal or entry of hepatocytes *in vitro*. Though 2F6 and RAM1 bind non-overlapping regions of the CSP-repeat domain, pretreatment with RAM1 abrogated 2F6’s capacity to block sporozoite traversal and invasion *in vitro*. Importantly, RAM1 reduced the efficacy of the polyclonal humoral response against CSP *in vivo,* paralleling the observed reduced efficacy of RTS,S in malaria-exposed populations. Taken together, our data demonstrate the interference of non-neutralizing antibodies with the efficacy of NAbs and may impact the efficacy of anti-CSP vaccines in malaria-exposed individuals.

## Introduction

*Plasmodium* parasites, the causative agents of malaria, are complex pathogens that have co-evolved with the human host. *Plasmodium* infection of mammals begins with injection of the sporozoite stage parasite into the skin by the bite of an infected *Anopheles* mosquito. The sporozoite surface is predominantly coated with the circumsporozoite protein (CSP). Antibodies against CSP are both the goal of the only licensed malaria vaccine, RTS/S AS01(Olotu et al., 2016; Stoute et al., 1997) and are found in exposed individuals within endemic areas(Tapchaisri et al., 1983).

Sporozoites traverse multiple cells in the skin prior to entering a capillary, which facilitates carriage of the sporozoite to the liver. Once in the liver, sporozoites traverse the sinusoidal endothelium, entering and exiting liver-resident macrophages and liver sinusoidal endothelial cells, and infect hepatocytes. Within hepatocytes, they develop as liver-stage (LS) parasites for the next 2-10 days(Shortt et al., 1948) without eliciting symptoms. This pre-erythrocytic stage of *Plasmodium* is a prime target for vaccine development as blocking the parasite at this stage would not only prevent disease progression, and related symptoms but also subsequent transmission thereby breaking the parasite life cycle. The development of pre-erythrocytic vaccines is particularly crucial for *Plasmodium* species *vivax* and *ovale*, which have unique latent forms of the parasite in the liver called hypnozoites that initiate relapse infections(Mueller et al., 2009). After extensive replication within the hepatocyte, parasites exit the liver and re-enter the bloodstream as merozoites and infect erythrocytes. During blood-stage infection, parasites undergo asexual cycles of replication, with each replicative cycle resulting in the destruction of an erythrocyte and the release of additional infectious merozoites. These destructive cycles cause all disease-related symptoms in the human host.

Virtually all pre-erythrocytic malaria vaccine candidates are tested initially in experimental clinical trials in malaria naïve westerners using controlled human malaria infection (CHMI). To date, two major approaches have been taken to elicit protection against *Plasmodium* sporozoite stages. The first approach uses a whole *P. falciparum* sporozoite (*Pf*SPZ)-based vaccine, where parasites are attenuated by irradiation, genetic manipulation or chemical inhibition. Parasites are then administered via mosquito bite(Herrington et al., 1991) or by intravenous administration of cryo-preserved whole sporozoites(Epstein et al., 2011; Hoffman et al., 2002) in CHMI settings(Epstein et al., 2011; Jongo et al., 2018; Sissoko et al., 2017). The other major approach is a subunit vaccine based on the major surface protein of the sporozoite, circumsporozoite protein (CSP). Several studies have shown that mAbs targeting the CSP repeat region can immobilize sporozoites upon pre-incubation (Hollingdale et al., 1984; Stewart et al., 1986) and lead to reduced infectivity when administered intravenously into passively immunized hosts(Charoenvit et al., 1991; Potocnjak et al., 1980). CSP-based vaccine development efforts have spanned multiple decades and culminated with the development of RTS,S/AS01 (Mosquirix™) by GlaxoSmithKline. RTS,S/AS01, a *Pf*CSP subunit vaccine formulation, elicits extremely high anti-CSP titers in non-immune westerners(Stoute et al., 1997) and malaria-endemic residents alike(Rts, 2015).

*Pf*SPZ and RTS,S/AS01 both elicit sterilizing immunity in non-immune subjects exposed to CHMI(Hoffman et al., 2002; Olotu et al., 2016). During its development, RTS,S/AS01 induced 50-87% sterile efficacy in CHMI (Gordon et al., 1995; Kester et al., 2008; Regules et al., 2016). However, in Phase III clinical trials of pediatric subjects in malaria endemic regions, RTS,S/AS01 did not induce sterile protection, but rather a ~30% partial protection was achieved(Olotu et al., 2016; Rts, 2015). The heterogeneity in CSP haplotype in the field had an effect on vaccine efficacy (VE) of RTS,S/AS01(Pringle et al., 2018), but it does not account for the complete lack of sterilizing immunity. Thus, achieving sterile efficacy against malaria infection in non-immune western adults does not consistently translate to protection in pre-immune subjects in malaria endemic areas. The mechanisms by which preexisting immunity reduces vaccine efficacy and the key factors influencing poor efficacy of vaccine-induced immunity are largely unknown, constituting a key roadblock to the development of an efficacious vaccine to eliminate malaria.

Previous work has demonstrated that antibodies contribute to protection from pre-erythrocytic infection(Aliprandini et al., 2018), and anti-CSP monoclonal antibodies (mAbs) are protective *in vivo*(Julien and Wardemann, 2019; Kisalu et al., 2018; Murugan et al., 2018; Tan et al., 2018; Triller et al., 2017). However, the B cell response against CSP is complex, and it is not yet fully known what antibody characteristics are critical to vaccine efficacy. Antibody maturation appears to be one key, as potent neutralization is linked with antibody affinity to CSP(Julien and Wardemann, 2019; Triller et al., 2017). Further, an epitope target at the R1/Repeat region boundary of CSP has been identified as a site of vulnerability(Kisalu et al., 2018). It has also been shown that anti-CSP antibodies are most effective in the skin, where they can render the sporozoite immobile thereby stopping the invasion process(Aliprandini et al., 2018). While these factors hint at what a desirable vaccine-elicited antibody response may need to look like, it also implies that much of the antibody response may be ineffective or irrelevant. Currently it is not known what effect these so-called irrelevant, non-neutralizing antibodies (nNAbs) have on vaccine efficacy or how their preexistence in endemic areas may interfere with vaccine-elicited immunity. Here, using the rodent malaria model *P. yoelii* 17XNL, we characterized how the presence of non-neutralizing antibodies impacts the efficacy of neutralizing anti-CSP vaccines. Our findings implicate direct antibody interference as a novel mechanism by which pre-existing nNAbs interfere with potent, vaccine elicited NAbs to reduce vaccine efficacy.

## Results

### Isolation and generation of RAM1

CSP is a complex protein with multiple unique domains and long stretches of repetitive motifs. The repeat regions consist of major and minor repeat units that can slightly differ within the molecule, and structurally appear to be spiral shaped with a specific chirality(Oyen et al., 2018). This unique structure enables an unusual binding profile to repeat antibodies, where it has been reported that up to eight antibody molecules (with a combined molecular mass of ~1200 kDa) can bind to a single molecule of CSP (approximately 70 kDa)(Lewis et al., 2020). Thus, CSP antibody binding and neutralization is complex, with the possibility of multiple types of antibodies interacting with a single molecule. 2F6 is a prototypical anti-CSP NAb(Sack et al., 2014), and binds the CSP repeat region. 2F6 confers protection in *in vitro* inhibition of sporozoite infection (ISI) experiments and in passive immunization experiments followed by mosquito-bite challenge *in vivo*(Harupa et al., 2014; Sack et al., 2014).

To characterize anti-CSP antibody-mediated protection in more depth, we sought to isolate new anti-CSP mAbs with properties that differ from the canonical NAb 2F6. We immunized BALB/cJ mice with recombinant *Py*CSP protein (*Py*CSP[NXA]), where NXA denotes that the *N-* linked glycosylation sequence (NXS/T) was mutated to NXA to prevent the post-translational addition of *N-*linked glycans. After multiple rounds of immunization, we sorted antigen-positive, class-switched memory B cells (MBCs) using *Py*CSP[NXA] as B cell tetramer bait (Fig. 1A). The IgHV and IgKV regions were amplified by RT-PCR and cloned into our expression backbone for recombinant antibody expression in HEK293 cells. The heavy and light chain genes of RAM1 are unique from that of 2F6. RAM1 utilizes the variable genes IgHV1-81*01 and IgKV8-30*01, whereas 2F6 utilizes IgHV9-2-1*01 and IgKV8-30*01.

**Fig. 1.**
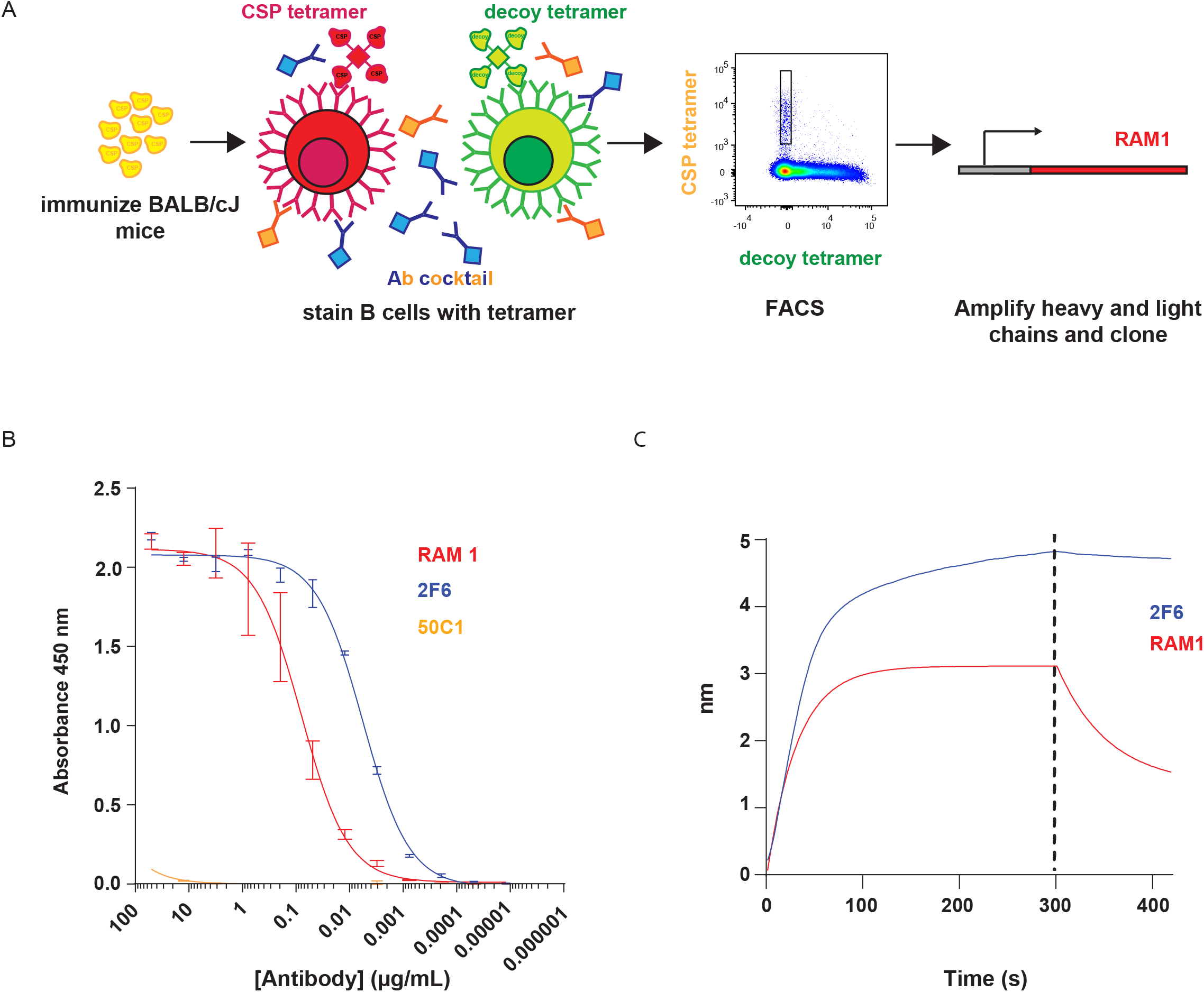
Generation and characterization of RAM1, a mAb against *P. yoelii* circumsporozoite protein. **(A)** Cartoon depicting the RAM1 production pipeline starting from immunization of BALB/cJ mice, staining of *Py*CSP[NXA]-specific B-cells using antigen and decoy tetramers by FACS followed by rescue and assembly of the heavy and light chains of RAM1 in pcDNA3.4 vector by RT-PCR and Gibson assembly reaction, respectively. **(B)** ELISA demonstrating the affinity of 2F6, RAM1 and 50C1 monoclonal antibodies to *Py*CSP[NXA]. **(C)** The real time binding affinity showing the association (0-300 sec) and dissociation (300-420 sec) phases of RAM1 and 2F6 to *Py*CSP[NXA], as determined by Octet, a Bio-Layer Interferometry (BLI) method. The results are representative of three independent experiments.

### RAM1 and 2F6 exhibit alternative binding kinetics to CSP

Both RAM1 and 2F6 bound to *Py*CSP[NXA] by ELISA, but with differing kinetics (Fig. 1B). RAM1 bound with an EC_50_ of 0.026 μg/ml, whereas 2F6 bound with an EC_50_ of 0.0093 μg/ml, indicating that 2F6 bound more tightly by an order of magnitude. The negative control mAb, 50C1, which binds to *Pf*TRAP with high affinity, exhibited no binding to *Py*CSP[NXA] and was used as a isotype control in subsequent experiments. We also assessed binding by real time kinetics using Octet, a Bio-Layer Interferometry method, enabling us to quantify on and off rates (Fig. 1C). 2F6 binds to *Py*CSP[NXA] with nanomolar affinity of 1.202×10^−9^ M in a 1:1 binding model, consisting of a fast k_on_ and a very slow k_off_ (Fig. 1C). Thus, 2F6 quickly associates and dissociates slowly, if at all. This high affinity kinetics profile is likely the genesis of its potent neutralizing activity, as affinity of anti-CSP mAbs is associated with neutralization in *P. falciparum*(Murugan et al., 2018). In contrast, when compared with 2F6, RAM1 binds to *Py*CSP[NXA] with more than a Log^10^ lower affinity (~2.66×10^−8^ M) (Fig. 1C). The binding profile does not fit well with a 1:1 binding model, indicating a higher stoichiometry of RAM1 binding CSP. RAM1 exhibits a relatively fast association rate (k_on_= −6×10^−4^ 1/Ms), but also has a fast dissociation rate (k_off_= −1.75×10^−3^ 1/s). Together, these data demonstrate that RAM1 associates relatively quickly but also dissociates quickly, resulting in a lower overall affinity than 2F6.

To further dissect the binding properties of these two mAbs, we created Fabs by papain digestion. The digested fragment antibodies (Fabs) contain only a single Fv fragment with one heavy and one light chain fragment linked by a disulfide bond, preventing any bivalent binding or avidity effects. 2F6.Fab bound *Py*CSP[NXA] with a Kd of 5.24×10^−7^ M (Fig. 2A). In contrast, RAM1.Fab bound weakly to *Py*CSP[NXA] (Fig. 2B), and as a result we were unable to derive a measurable Kd. This suggests that the binding of RAM1 may be dependent on avidity effects. Thus, 2F6 stably binds to *Py*CSP[NXA] with high affinity in a kinetic model that fits a 1:1 binding ratio, which was confirmed by kinetic measurements as a Fab. In contrast, RAM1 binding did not fit a 1:1 binding model, and when evaluated as a Fab lost almost all of its reactivity with *Py*CSP[NXA]. Taken together, these data indicate that 2F6 and RAM1 recognize *Py*CSP[NXA] fundamentally differently, both in basic affinity and in potential binding modality. As these studies were conducted *in vitro* with recombinant *Py*CSP[NXA], we next evaluated 2F6 and RAM1 binding to *Py*CSP in its native form on the surface of the sporozoite. We stained *P. yoelii* sporozoites with 2F6 conjugated to AlexaFluor 405 and RAM1 conjugated with AlexaFluor 594. AlexaFluor-488 conjugate of 50C1 was used as a control. As expected, 50C1 did not bind to *P. yoelii* sporozoites. In contrast, RAM1 and 2F6 exhibited circumferential binding to *P. yoelii* sporozoites, consistent with binding to CSP (Fig. 2C).

**Fig. 2.**
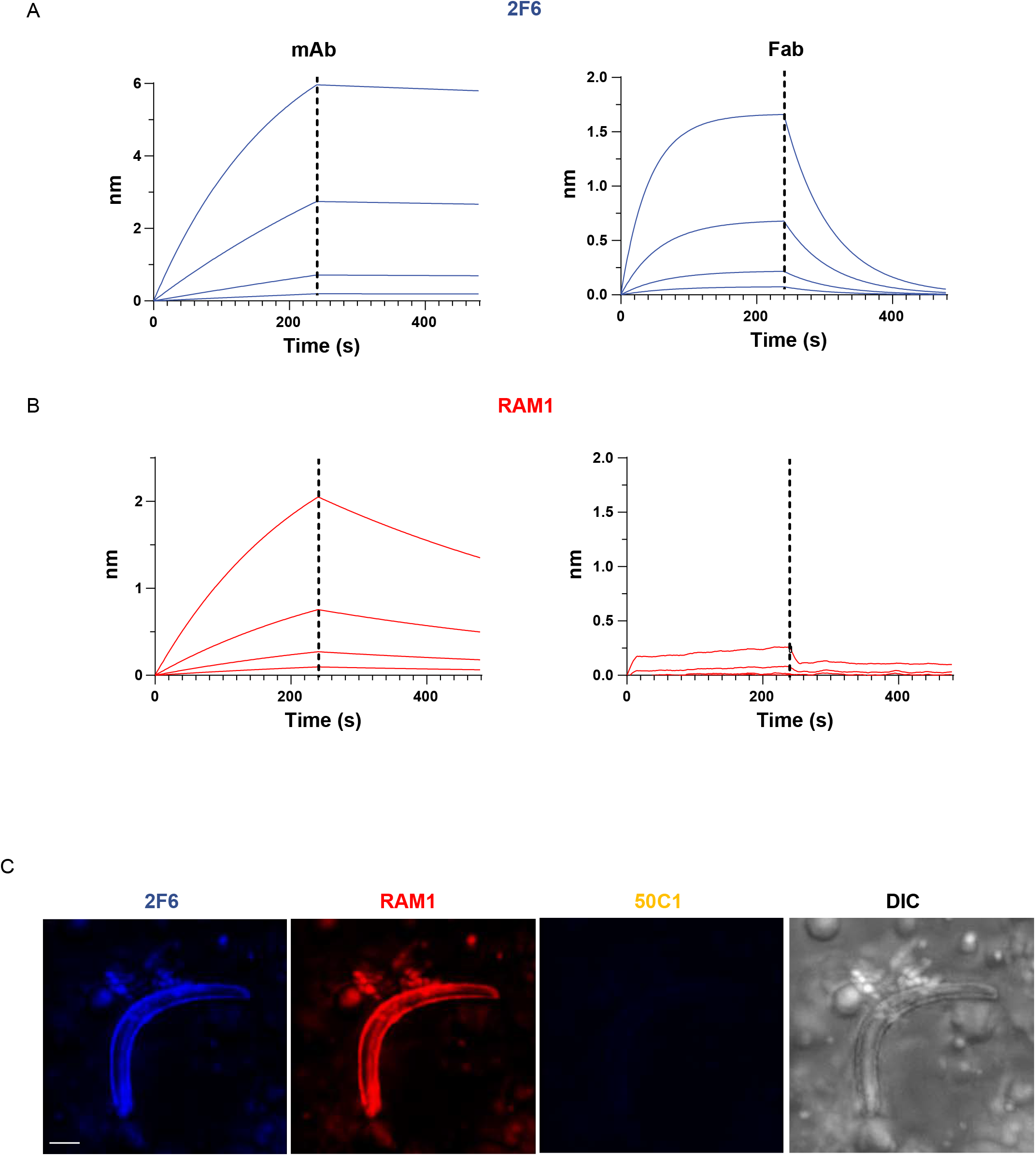
RAM1 binds to *Py*CSP[NXA] and *P. yoelii* sporozoites. **(A and B)** The association (0-240 sec) and dissociation (240-480 sec) kinetics of 2F6 (blue, A) and RAM1 (red, B) mAbs (left) and Fabs (right) to *Py*CSP[NXA] as determined by Octet. **(C)** Representative images of immunofluorescence assay (IFA) demonstrating the binding capacity of fluorescently labelled RAM1 (red), 2F6 (Blue) and 50C1 (control) monoclonal antibodies to *P. yoelii* sporozoites. Sporozoite morphology is shown in a differential interference contrast (DIC) image. Scale bar 5 μm. The results are representative of three independent experiments.

### RAM1 and 2F6 recognize non-overlapping epitopes within the central repeat domain of CSP

*Py*CSP is a GPI-anchored protein consisting of several different domains that are thought to have functionality at different points in the parasite life cycle(Swearingen et al., 2016). The protein is organized into domains consisting of the N terminal region, the R1 conserved junctional region, a central repeat domain, the R2 region, and the C-terminal region with highly conserved Thrombospondin type 1 repeats (TSR). To identify the binding sites of RAM1 and 2F6, we created domain deletion constructs lacking N-terminal *Py*CSP[ΔN] and C-terminal domains *Py*CSP[ΔC]. A third domain mutant was created that contains both the N-terminal and C-terminal domain, connected by two major repeat units *Py*CSP[N2rC] (Fig. 3A). We tested the binding efficiency of the mAbs RAM1, 2F6 and 50C1 to all domain variants (Fig. 3B). Both 2F6 and RAM1 bound to full-length *Py*CSP, *Py*CSP[ΔN] and *Py*CSP[ΔC] constructs, suggesting that both mAbs bind to an epitope within the central repeat region that is common to each construct. RAM1 also bound to *Py*CSP[N2rC], which contains two major repeat motifs, whereas 2F6 did not (Fig. 3B). Thus, both mAbs bind the repeat regions, but appear to have different specific epitopes.

**Fig. 3.**
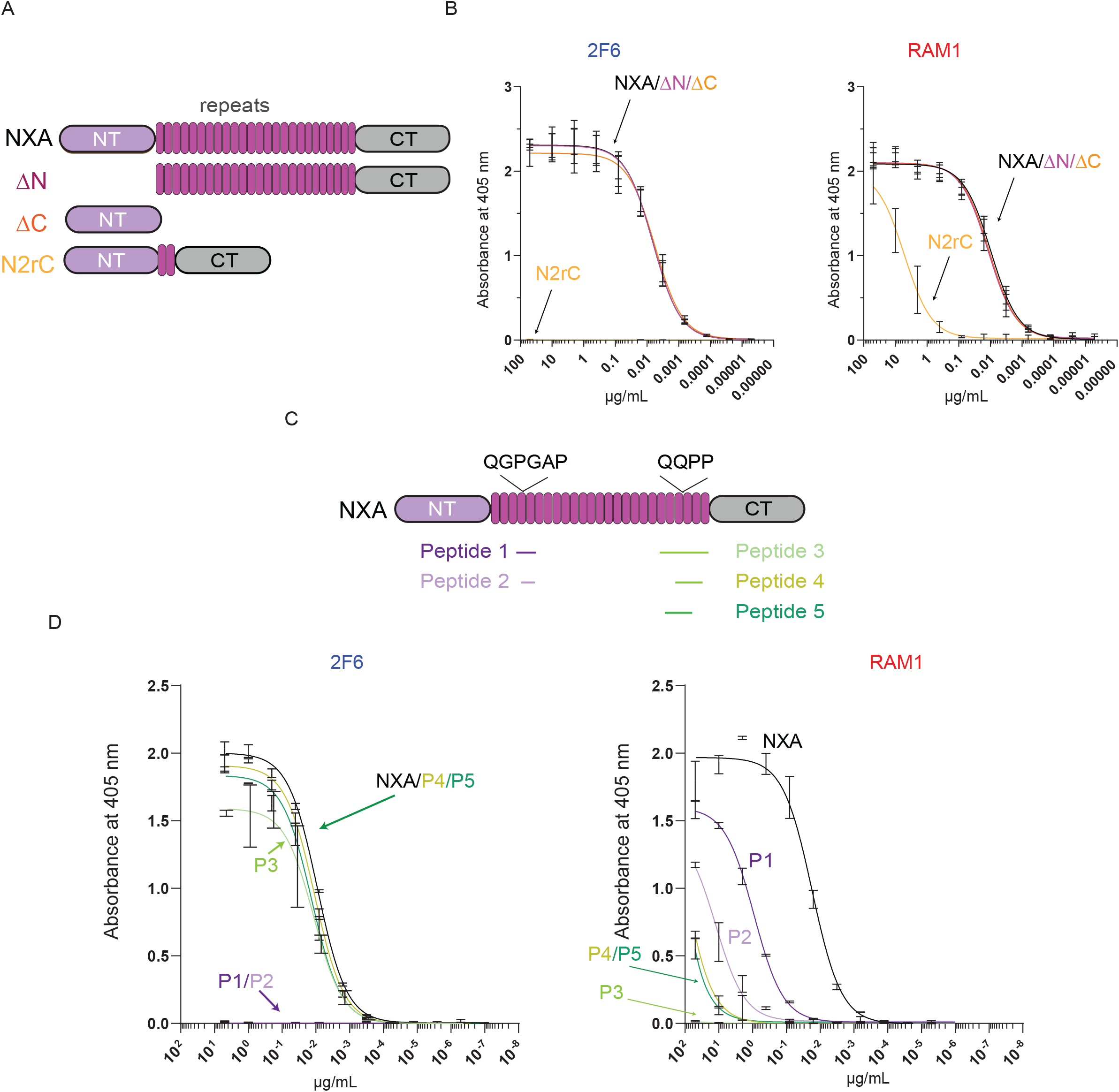
RAM1 binds to the major repeat of *Py*CSP[NXA]. **(A)** Schematic representing *Py*CSP[NXA] constructs used; full length (NXA), devoid of N-terminal domain (ΔN), devoid of C-terminal domain (ΔC) and N, C-terminal domains connected with 2 major repeat units (N2rC), used in this study. The N-terminal (NT), C-terminal (CT) and the repeats are shown in violet, grey and purple, respectively. **(B)** ELISA demonstrating the binding of 2F6 and RAM1 to *Py*CSP[NXA] constructs depicted in (**A**). **(C)** Schematic diagram representing the location of synthesized peptides representing the central repeat region of *Py*CSP[NXA] used in this study **(D)** The binding affinity of RAM1 and 2F6 to the different *Py*CSP[NXA] repeat region peptides as demonstrated by ELISA. The results are representative of three independent experiments.

*Py*CSP has a large number of repeat units consisting of two motifs: the proximal major repeat (QGPGAP) and distal minor repeat (QQPP) (Fig. 3C). This arrangement is analogous to *Pf*CSP, which also has major and minor repeat motifs, although the specific amino acid sequences are different(Coppi et al., 2005). To map the specific binding of RAM1 and 2F6 within the repeat motifs, we synthesized a series of peptides that spanned different regions within the *Py*CSP repeat region (Fig. 3C). 2F6 bound peptides spanning the minor repeat (QQPP) but did not bind to any of the peptides spanning the major repeats (Fig. 3D). 2F6 binding to minor repeat peptides was similar in magnitude to that of full length *Py*CSP[NXA], indicating that the entire epitope was likely captured by minor repeat peptides. In contrast, RAM1 bound peptides spanning the proximal major repeat region (QGPGAP) and had no reactivity to peptides spanning the minor repeat region (Fig. 3D). These results are consistent with the finding that RAM1, but not 2F6, binds *Py*CSP[N2rC], which contains two major repeat units and no minor repeats. Interestingly, RAM1 bound major repeat peptides with a lower affinity than the binding to *Py*CSP[NXA]. Thus, either the epitope is more complex than can be presented by the peptide, or a subset of the interactions that mediate RAM1 binding could not be replicated by a linear peptide. Regardless, these findings illustrate that RAM1 and 2F6 recognize different repeat regions within *Py*CSP[NXA], and that their epitopes are non-overlapping.

### RAM1 does not neutralize *P. yoelii* sporozoite infection

After deposition within the skin during the probing of an *Anopheles* mosquito, sporozoites must leave the dermis and enter a capillary to survive and further develop. To do this, sporozoites make use of their specialized capacity to traverse cells using a perforin-like mechanism (Ishino et al., 2005). While the sporozoite can traverse any cell type in a cell-type-independent manner, the dermis is enriched for fibroblasts. Thus, we chose to assess the fibroblast traversal ability of sporozoites in the presence of RAM1. Specifically, we incubated freshly isolated *P. yoelii* wild type sporozoites with different concentrations (1, 5 and 10 μg) of RAM1, 2F6 or 50C1 antibodies for 10 mins and allowed sporozoites to traverse human foreskin fibroblast HFF-1 cells for 90 mins in the presence of high molecular mass dextran (70 kDa). The dextran positive cells are wounded cells and were quantified using flow cytometry. We observed no significant differences between the traversal rate of fibroblasts in the presence of RAM1. As expected, we observed decrease in traversal in the presence of 2F6, a known anti-CSP NAb (Fig. 4A). Next, we asked whether RAM1 interferes with infection of hepatocytes. We infected Hepa 1-6 cells for 90 mins with *P. yoelii* sporozoites pre-incubated for 10 mins with different concentrations (1, 5 and 10 μg) of RAM1, 2F6 or 50C1 (Fig. 4B). We observed no significant differences in the invasion of hepatocytes by RAM1 or 50C1 pretreated sporozoites. In contrast, we observed a dose-dependent decrease in hepatocyte entry in the presence of 2F6. This suggests unlike 2F6, which blocks both sporozoite traversal and invasion, RAM1’s activity is consistent with that of a nNAb.

**Fig. 4.**
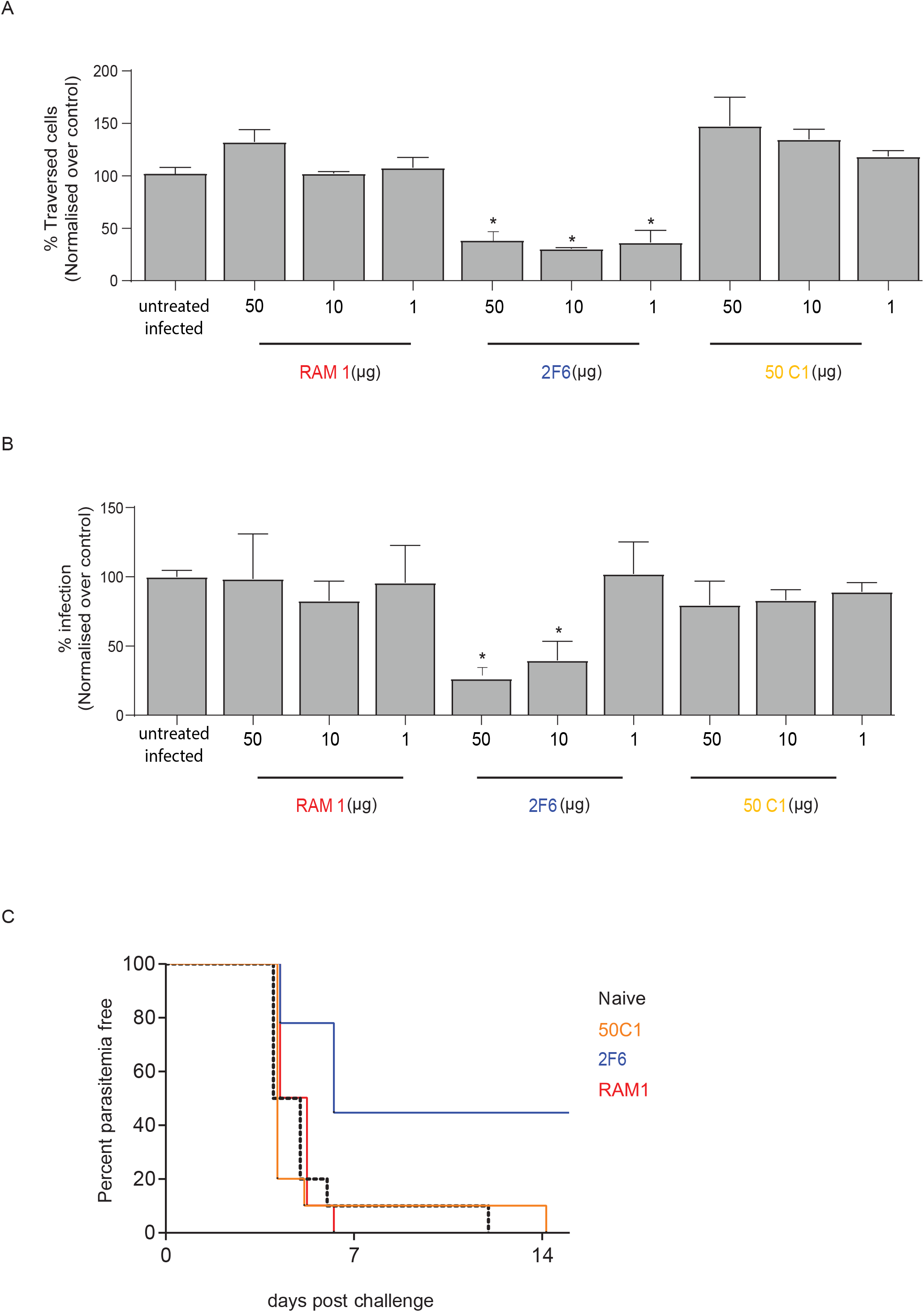
RAM1 does not neutralize sporozoite activity *in vitro* or *in vivo*. Freshly isolated sporozoites were pre-incubated with different concentrations of RAM1, 2F6 and 50C1 antibodies for 10 mins. **(A)** Antibody-treated sporozoites were exposed to HFF-1 cells in the presence of Dextran-FITC for 30 mins. The bar graph represents the percentage of cells traversed as assessed by dextran positive cells. n=3; * denotes p<0.05. **(B)** Hepa 1-6 cells were infected with antibody-treated sporozoites for 90 min to assess hepatocyte entry. The bar graph here represents the percentage of hepatocytes that were CSP-positive as evaluated by flow cytometry. n=3; * denotes p<0.05. **(C)** BALB/cJ were injected with 150 μg of RAM1, 2F6 or 50C1 intraperitoneally. After 24 h, mice were challenged with bites from *P. yoelii* infected mosquitoes and patency was assessed from day 4 through day 14. The graph represents percentage parasitemia free mice over time, including 10 mice from 2 independent experiments.

We next assessed the capacity of RAM1 to alter sporozoite infectivity *in vivo*. BALB/cJ mice were injected intraperitoneally with 150 μg of RAM1, 50C1 or 2F6. After 24 h, mice were challenged with *P. yoelii* infected mosquitoes. Patency was evaluated beginning four days after infection and continued through fourteen days post-challenge. Nearly all naïve and 50C1 treated mice were infected within 6 days of challenge (Fig. 4C, Supplementary Table 1). Consistent with the observations of others(Sack et al., 2014; Tarun et al., 2007), we observed 50% protection in mice that were passively infused with 2F6. Consistent with the *in vitro* results, RAM1 treatment resulted in 100% infection within 6 days of infection, again suggesting that RAM1 is a nNAb. Taken together, these data demonstrate that RAM1 binds *P. yoelii* CSP but does not impact its capacity to traverse or infect cells *in vitro* or *in vivo*.

### RAM1 reduces the inhibitory activity of 2F6 *in vitro*

We next asked if RAM1 could impact the anti-parasitic activity of the anti-CSP NAb 2F6. We pre-incubated *P. yoelii* sporozoites with pairwise combinations of 2F6, RAM1 and the non-*P. yoelii* binding negative control mAb 50C1. We then assessed the traversal of HFF-1 fibroblasts (Fig. 5A). Interestingly, the concurrent exposure of sporozoites to 2F6 and 50C1 significantly inhibited sporozoite traversal. In contrast, concurrently treating sporozoites with a mixture containing equal amounts of RAM1 and 2F6, or a 5:1 excess of RAM1, resulted in traversal rates similar to untreated sporozoites (Fig. 5A). This suggests that the presence of RAM1, whether in equal amounts or in excess, was sufficient to significantly reduce 2F6’s capacity to inhibit sporozoite traversal.

**Fig. 5.**
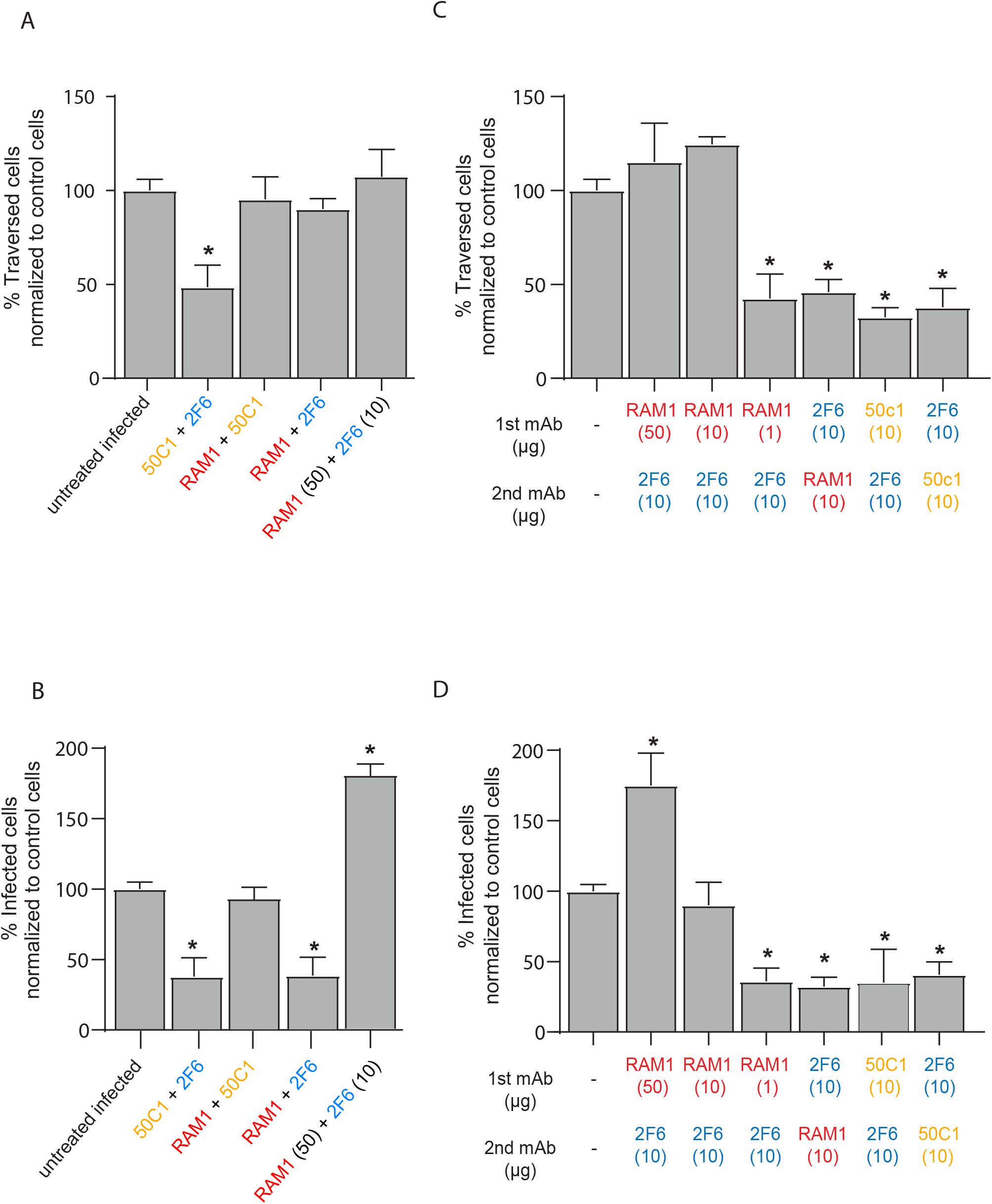
RAM1 potentiates the blocking activity of an anti-CSP NAb. Freshly isolated *P. yoelii* sporozoites were pre-incubated with pairwise combinations of RAM1, 2F6 and 50C1 antibodies concurrently (A & B) or sequentially (C & D) for 10 min each. Antibody treated sporozoites were exposed to HFF-1 cells in the presence of Dextran-FITC for 30 mins to assess traversal capacity (A & C) or Hepa1-6 cells for 90 mins to assess hepatocyte entry by sporozoites (B & D). n=3; * denotes p<0.05.

In contrast, the presence of RAM1 in equal amounts did not impact the hepatocyte invasion inhibitory activity of 2F6 (5B). Concurrent treatment of sporozoites with equal amounts of RAM1 and 2F6 exhibited similar levels of hepatocyte invasion, compared to 2F6-50C1 combination treatment. Consistent with traversal, concurrent treatment of RAM1 and 50C1 exhibited levels of hepatocyte entry similar to uninhibited sporozoites, reflecting RAM1’s non-neutralizing activity (Fig. 5B). However, when RAM1 was added concurrently in a 5:1 excess, it completely inhibited the invasion blocking activity of 2F6. Thus, the presence of RAM1 ablated the anti-traversal activity of 2F6, whereas hepatocyte invasion blocking was only ablated when RAM1 was present in excess. These findings imply that vaccine-elicited anti-CSP mAbs could be ineffective against hepatocyte infection if non-neutralizing antibodies were also present, as seen in pre-existing immunity, particularly if they are present in excess of any vaccine-elicited NAbs. In the skin, where traversal is thought to play a critical role in protection, the presence of nNAbs in even equal amounts appears to be particularly problematic.

To further dissect the interference phenotype of RAM1 observed above, we sequentially preincubated sporozoites with combinations of RAM1, 2F6 and 50C1 for 10 mins each and then assessed traversal of HFF-1 fibroblasts (Fig. 5C) and infection of Hepa 1-6 cells (Fig. 5D). Pretreatment of sporozoites with 2F6 followed by treatment of RAM1 or 50C1 reduced sporozoite traversal of fibroblasts (Fig. 5C) or entry of hepatocytes (Fig. 5D) at similar levels to 2F6 alone. In contrast, pretreatment with RAM1 (10 or 50 μg), followed by 2F6 (10 μg) led to a significant impairment in the capacity of 2F6 to inhibit fibroblast traversal or hepatocyte invasion (Fig. 5C, D). Thus, if present prior to 2F6, RAM1 was able to completely prevent 2F6 from neutralizing the sporozoite. While sequential exposure of antibodies is not likely to occur in this manner *in vivo*, these findings highlight a complication for vaccination, where nnAbs can interact with the antigen prior to exposure to potential NAb B cell receptor precursors. In this scenario, our results imply that such BCRs may be prevented from interacting with the antigen when exposed to pre-existing nnAbs after injection.

To further understand the functional consequence of these antibody combinations on the surface of the sporozoite, we evaluated the extent of the circumsporozoite precipitation reaction (CSPR). CSPR is a well characterized phenomena where CSP is secreted at the apical end of sporozoites and then translocated backwards, forming a thread-like precipitate at the posterior end of the sporozoite in the presence of anti-CSP NAbs(Stewart and Vanderberg, 1991). Incubation of sporozoites with 2F6 and RAM1, either concurrently or by first adding 2F6 first resulted in large, well-delineated extensions of CSPR (Fig. 6A, B). In contrast, when RAM1 is added first, we observed a far-reduced rate of CSPR (Fig. 6A, B). Interestingly, in sporozoites treated first with RAM1, we observed a different phenomenon. These sporozoite exhibit vesicles that appear to be secreted that measure approximately 300 nm in diameter and are CSP positive. This phenomenon is far less prevalent in sporozoites that have been exposed to 2F6 first or concurrently (Fig. 6A, C). Together these data suggest that RAM1 exposure induces an altered state of the sporozoite, potentially rendering it less susceptible to recognition or neutralization by NAbs. As noted above, this relationship could have profound functional consequences for vaccination with attenuated sporozoites in the context of pre-existing nnAbs.

**Fig. 6.**
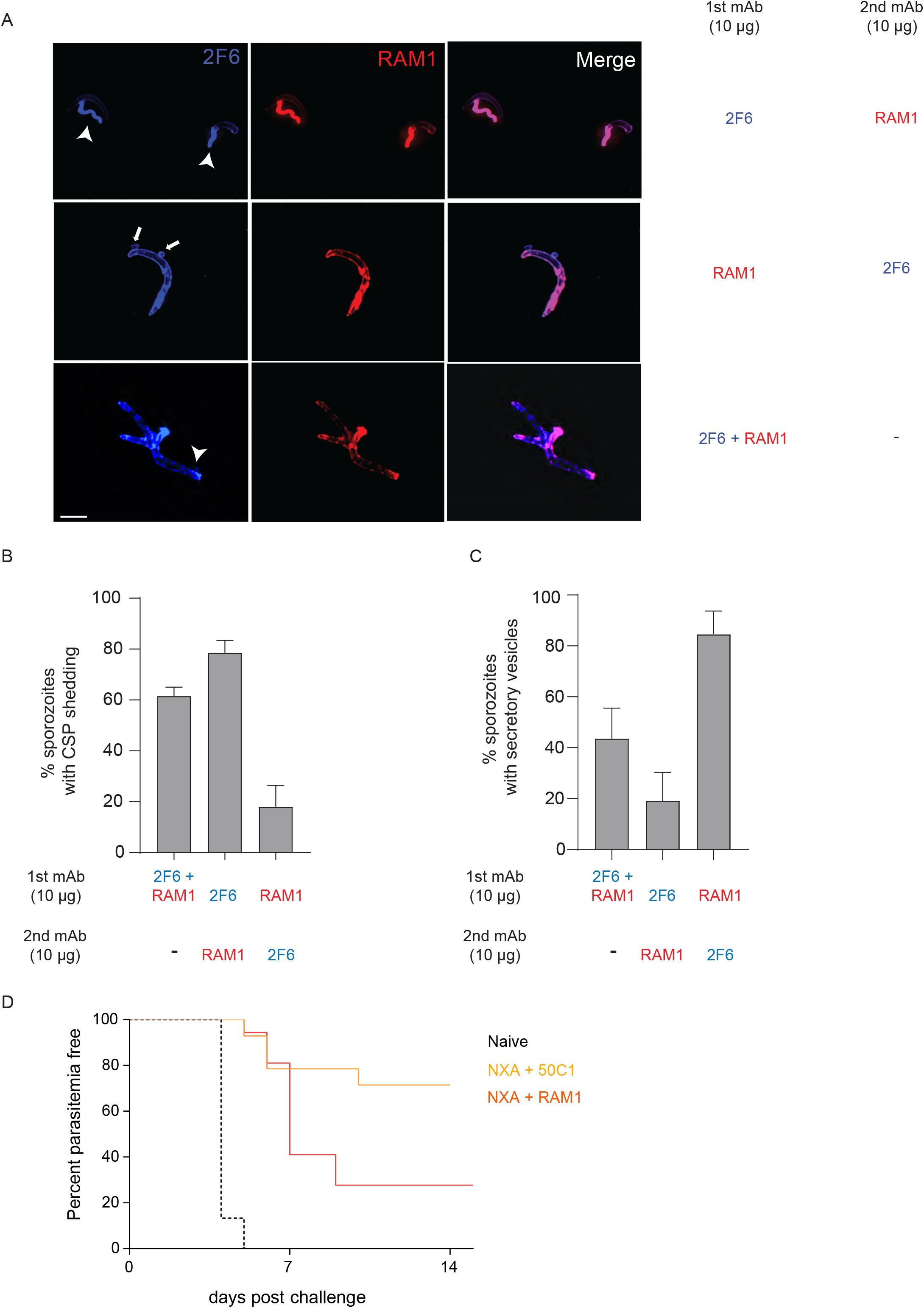
RAM1 alters protective immunity induced by a CSP-directed recombinant vaccine. **(A)** Immunofluorescence of *P. yoelii* sporozoites incubated with 2F6-Alexa 405 (blue) and RAM1-Alexa 594 (red) together or sequentially for 10 min each. The arrowhead points to the regions of CSP reaction or shedding (CSPR). The full arrows point to vesicles observed, which appear to bud from the sporozoites. Scale bar 5 μm. **(B)** The bar graph represents the percentage of sporozoites that exhibit CSPR. n=3; * denotes p<0.05. **(C)** The bar graph represents the percentage of sporozoites that exhibit defined vesicles. n=3; * denotes p<0.05. **(D)** BALB/cJ were immunized with *Py*CSP[NXA] at 0, 2 and 6 weeks followed by passive transfer of 500 μg of RAM1, 2F6 and 50C1 antibodies intraperitoneally. After 24 h, mice were challenged with *P. yoelii* infected mosquitoes and patency were assessed from day 3 through day 14. The graph represents percentage parasitemia free mice over time. Data are from 14-15 mice across 3 independent experiments.

### RAM1 reduces protective immunity induced by a subunit CSP vaccine

To alter the outcome of individuals in malaria-endemic areas that have been immunized with a malaria vaccine, pre-existing nNAbs would need to both be effective at impeding the activity of a NAb *in vivo* and would need to diminish the potency of a vaccine-induced polyclonal response. To evaluate whether RAM1 could interfere in the *in vivo* setting of CSP immunization, live infection challenge, we immunized BALB/cJ mice with recombinant *Py*CSP[NXA] for 0, 2 and 6 weeks. After final immunization, 500 μg 50C1 or RAM1 was passively transferred to mice to mimic an excess of nNAbs at the time of challenge. Twenty-four hours following passive transfer, mice were challenged by the bite of *P. yoelii*-infected *Anopheles* mosquitoes. Patency was assessed daily beginning three days post challenge. The immunization regimen resulted in ~70% sterile protection in mice that received the non-binding 50C1 control mAb passive transfer. Strikingly, the passive transfer of RAM1 into vaccinated animals resulted in protection of only ~25% of *Py*CSP[NXA] immunized mice (Fig. 6D, supplementary Table 2). Taken together, these findings suggest that RAM1 can interfere with the protection afforded by monoclonal NAbs *in vitro*, but it can also drastically diminish the levels of protection mediated by the polyclonal anti-CSP response elicited by vaccination *in vivo* in a setting similar to vaccination in the field.

## Discussion

Preexisting humoral immunity is the sum total of many different antibodies, with genetic makeup, epitope specificity, and binding characteristics all contributing to the specific features of the interaction between vaccine-elicited antibodies and their pathogen target. It was suggested decades ago that monoclonal antibodies had a greater capacity to block infection than polyclonal responses in mice(Charoenvit et al., 1991), consistent with the field observation that individuals with naturally occurring polyclonal responses to CSP are not reliably protected from subsequent infection(Offeddu et al., 2012). While it is tempting to characterize antibodies as simply ‘neutralizing’ or ‘non-neutralizing’, embedded within this portrayal is the assumption that antibodies act independently. Rather, our data suggest that the activity of neutralizing antibodies can be altered by antibodies that, on their own, neither boost nor inhibit infection.

Interestingly, the capacity of RAM1 to reduce 2F6’s neutralizing activity cannot be explained by a simple assessment of affinity or specificity. While 2F6 binds to *Py*CSP[NXA] with nanomolar affinity of 1.202×10^−9^ M (Fig. 1C, and 2B), RAM1 binds to *Py*CSP[NXA] with less affinity (~2.66×10^−8^ M) (Fig. 1C and 2A). Moreover, RAM1 binds the *Py*CSP[NXA] major repeat region (QGPGAP), whereas 2F6 binds the *Py*CSP[NXA] minor repeat region (QQPP) (Fig. 3D). From these data alone, we would not predict that RAM1 could outcompete the binding or functionality of 2F6.

However, a more detailed evaluation of the binding of each antibody suggests the potential for a complex interplay between the two molecules on the sporozoite surface, which contains a dense array of *Py*CSP molecules including many major and minor repeats. Our data suggest that 2F6 likely associates with *Py*CSP in a 1:1 binding model, consisting of fast k_on_ and very slow k_off_ rates. In contrast, RAM1 also exhibits fast k_on_ binding kinetics, but a much faster k_off_ than 2F6, consistent with a higher stoichiometry of binding. When antibodies were tested for binding as Fabs, RAM1 binding was dramatically reduced, consistent with the hypothesis that avidity is the major driver of RAM1 binding and stability. We predict that these disparate modes of binding facilitate competition between multiple RAM1 molecules and a single 2F6 molecule on the surface of the sporozoite. Indeed, when we expose *Py* sporozoites to both antibodies, we observe binding of both antibodies to sporozoites rather than saturation of 2F6 (Fig. 2C). Importantly, this competition also has functional consequences: when added prior to 2F6, or in higher concentration, RAM1 can abrogate the neutralizing function of the NAb (Fig. 5). Thus, avid binding of RAM1 to surface CSP (whether inter- or intra-molecular) appears to interfere with the high affinity binding and neutralization of 2F6, despite RAM1’s significantly lower affinity for CSP.

While our experiments were performed in a laboratory setting, it is tempting to speculate how inter-antibody interactions may impact vaccine efficacy in the field. Individuals in malaria-endemic areas have anti-CSP antibodies, originating from anti-CSP B cells, that do not achieve sterile protection from infection(Hoffman et al., 2002; Roestenberg et al., 2009). These anti-CSP B cells receive continual re-stimulation by natural exposure and are likely preferentially boosted by the administration of a CSP-based vaccine like RTS,S/AS01. It stands to reason that this process produces a robust population of anti-CSP nNAbs. Even if this occurs in parallel with the priming and boosting of new protective antibodies during vaccination, the polyclonal response will be a compilation of antibodies, some which may potentiate the activity of others. While it would be a reasonable hypothesis that antibodies that shared binding epitopes may have the greatest likelihood of competing, our data suggest that the nNAb RAM1, which does not share an epitope with the NAb 2F6, can still diminish the neutralization capacity of the NAb.

Direct interference between nNAbs and NAbs at the sporozoite surface represent only one of many challenges that exist to vaccination in the context of preexisting immunity. In addition to playing a role at the time of sporozoite challenge, which we explore here, nNAbs could also directly bind the vaccine antigen after injection but prior to B cell recognition. Our findings imply that in cases like RAM1, this initial interaction may be able to prevent the NAb B cell precursor from recognizing CSP and becoming stimulated, preventing the generation of robust NAbs. Further, nNAbs may represent original antigenic sin, where nNAb memory B cells are preferentially utilized by the immune system in response to vaccination. Thus, new potential NAb precursor B cells may be at a competitive disadvantage, resulting in a reduced ability to induce potent NAbs. This may also result in a stoichiometric imbalance where pre-existing nNAbs circulate in relative abundance compared to NAbs in response to vaccination. As shown in this study, this stoichiometric imbalance can directly lead to reduced efficacy. Thus, deciphering the immunological consequences of pre-existing immunity on vaccine efficacy remains an important area for future investigation.

Despite extensive vector control efforts, highly effective antimalarial drugs and several vaccine candidates that have shown promise in malaria-naïve individuals, malaria remains a global health problem worldwide. It is widely appreciated that a vaccine that is highly effective in malaria-endemic populations would contribute dramatically to malaria eradication efforts. Yet, malaria vaccine development efforts continue to suffer from a severe disconnect between promising vaccine efficacy in malaria-naïve westerners and a lack of sterilizing vaccine efficacy in endemic regions. It is now well-recognized that pre-existing immunity to malaria is a key roadblock to the development of an effective vaccine(Good, 1995). Our work demonstrates that antibody interference between nNAbs and NAbs may contribute to this lack of translatability in endemic populations. A better understanding of the mechanisms by which vaccines fail to induce robust protection in malaria-endemic areas could inform next generation vaccines and help to control malaria world-wide.

## Materials and Methods

### Recombinant proteins, deletion constructs and peptides

Amino acids 21-362 of *Plasmodium yoelii* <u>C</u>ircum<u>s</u>porozoite <u>P</u>rotein (PlasmoDB:PYYP_0405600) (*Py*CSP) was cloned into a pcDNA3.4 mammalian expression vector (Thermo Fisher, Waltham, MA, USA) containing a CMV immediate early promoter, a tPA signal, and a C-terminal 8×HIS tag, followed by an AviTag. The amino acids at positions A and B of *Py*CSP were mutated to an Alanine to remove the predicted *N-*glycosylation sites on the protein (this construct is termed *Py*CSP[NXA]). Protein construct, *Py*CSP[NXA] was produced in suspension HEK293 cell cultures, as previously described(Harupa et al., 2014; Kaushansky et al., 2015). Briefly, plasmid DNA was transfected into HEK293 cells using PEI MAX 40000 (Polysciences, Inc., Warrington, PA, USA). The cultures were grown in FreeStyle 293 serum-free media (Thermo Fisher) for five days and then the supernatant containing the protein was harvested by pelleting cells at 4000 rpm for 20 mins at 4°C. The supernatant was adjusted with 350 mM NaCl and 0.02% sodium azide and the protein in the supernatant was purified by Ni-affinity and size exclusion chromatography. The protein was eluted and stored in 10 mM HEPES (pH 7.0), 150 mM NaCl, 2 mM EDTA at 4°C for short-term use or at −20°C for long-term. For capture ELISAs and fluorescence activated cell sorting (FACS) experiments, biotinylated proteins were used. Biotinylation of AviTag-containing *Py*CSP[NXA] was performed using a BirA biotin-protein ligase kit (Avidity, LLC; Aurora, CO, USA), according to manufacturer’s instructions. Post-biotinylation, proteins were purified by size-exclusion chromatography and concentrated.

For epitope mapping studies, the genes encoding the full-length *Py*CSP[NXA] gene devoid of N-terminal region, *Py*CSP[ΔN], the C-terminal region *Py*CSP[ΔC], and most of the central repeat region except for two major repeat units *Py*CSP[N2rC] were cloned in pcDNA3.4 vector and produced in HEK293 suspension cultures. These deletion constructs were expressed and purified as described before^41^. N-terminal biotinylated peptides of *Py*CSP[NXA] listed in Table 1 were obtained from GenScript and used in epitope mapping ELISAs.

**Table 1.**
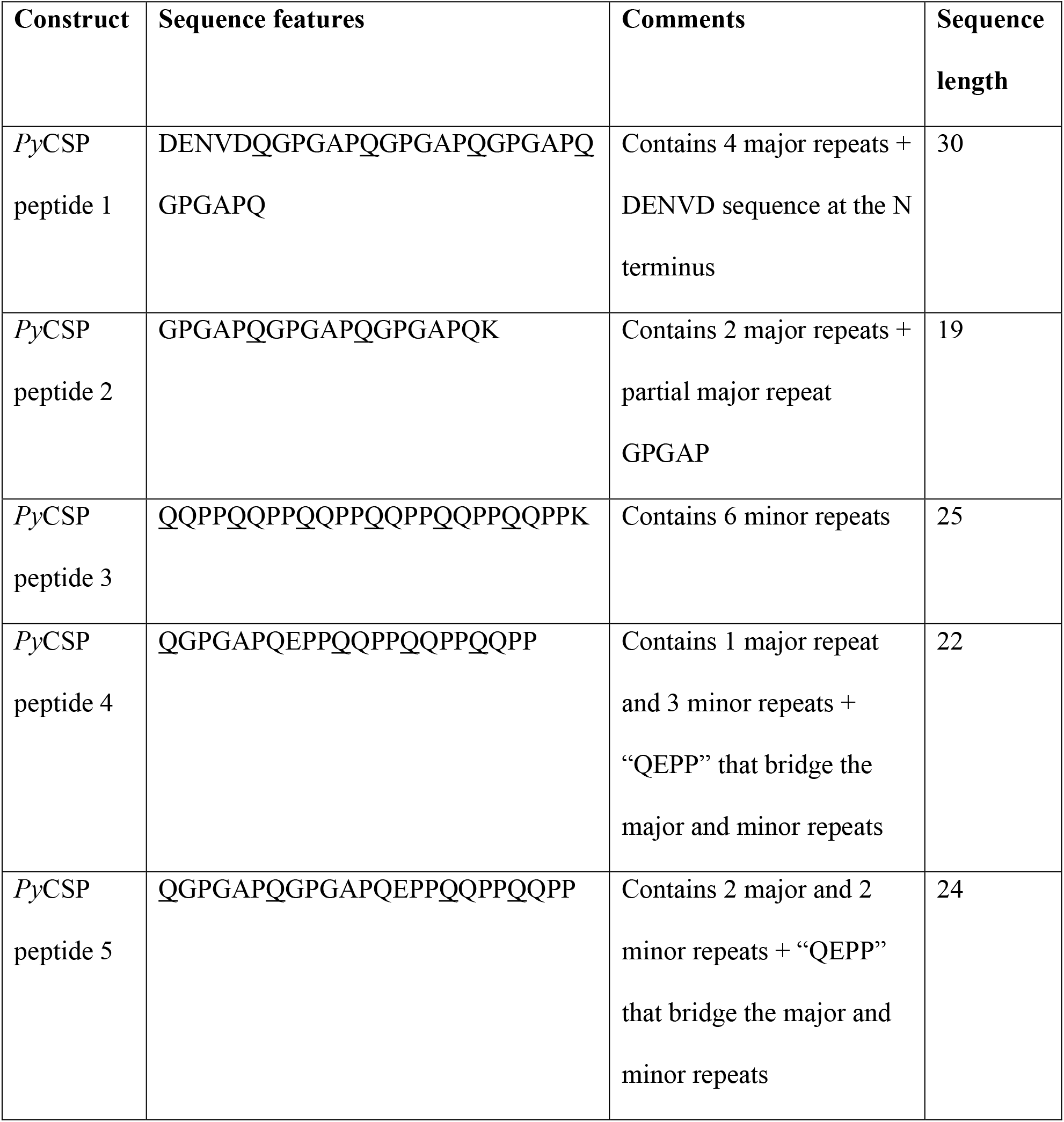
List of *Py*CSP peptides used in the study. Related to Figure 3.

### Antigen-specific B cell sorting and cloning of monoclonal antibodies

Female BALB/cJ mice, 6-8 weeks of age, were immunized with 20 μg of *Py*CSP[NXA] formulated with 20% v/v of Adjuplex (Invivogen, San Diego, CA, USA) at 0, 2 and 6 weeks. At week 7, spleens were harvested and splenocytes were isolated by homogenizing the spleen tissue and passing the cells through a 70 micron cell strainer. Following splenocyte isolation, B cells were enriched by negative selection using EasySep mouse B cell enrichment kit as per manufacturer’s instructions (StemCell Technologies Inc., Tukwila, WA, USA). Enriched B cells were then resuspended in FACS buffer and stained and sorted as described previously(Carbonetti et al., 2017). Briefly, B cells were incubated with anti-mouse CD16/CD32 (mouse Fc block; BD Biosciences) and a decoy tetramer (BV510) for 10 mins at room temperature. Following this, cells were stained with B220-PacBlue, CD38-APC, IgM-FITC, IgD-AF700 (BioLegend, San Diego, CA, USA) and *Py*CSP[NXA] tetramer (BV786) for 30 mins at 4°C. Cells were then washed and resuspended in FACS buffer, and passed through a 30 micron filter before sorting on a BD FACS Aria II with a 100-μm nozzle running at 20 psi. *Py*CSP[NXA]-specific class switched memory B cells (B220^+^ CD38^+^ IgM^−^IgD^−^antigen^+^ decoy^−^cells) were sorted at 1 cell per well into 96-well PCR plates and stored at −80°C until use. cDNA synthesis and amplification of the IgG and IgK variable regions from sorted cells were performed as previously described(von Boehmer et al., 2016). Heavy and light chain PCR amplicons were cloned into pcDNA3.4 expression vectors containing murine IgG1and IgK constant regions, respectively, by Gibson assembly.

### Expression and purification of recombinant mAbs

Plasmid DNA encoding the heavy and light chains of recombinant monoclonal antibodies and polyethylene imine (Polysciences, Warrington, PA) were mixed at a 1:4 ratio, respectively, and co-incubated in sterile PBS for 15 mins at room temperature. Following incubation, this mixture was applied to mammalian FreeStyle™ HEK293-F cell culture (EMD-Millipore, Darmstadt, Germany) at a density of 1 million cells per milliliter. Cultures were placed at 37°C, 5% CO^2^ on a shaker platform for 5 days and then cells were removed from the supernatant by centrifugation at 4000 rpm for 20 mins. The clarified supernatant was then passed over a HiTRAP MabSelect Sure column (GE Lifesciences #11003493) and eluted from the column per the manufacturer’s protocol. The eluate was subsequently buffer exchanged into HBS-E (10 mM HEPES, pH 7, 150 mM NaCl, 2 mM EDTA) and the size and purity of each antibody were determined by size exclusion chromatography using a Superdex 200 10/300 column (GE Healthcare, Chicago, IL, USA).

### RAM1 and 2F6 Epitope Mapping ELISA

#### Deletion Constructs

RAM1 and 2F6 epitope mapping against *Py*CSP deletion constructs was performed by direct ELISA. All incubations were done at 37°C for 1 h, unless stated otherwise. Immulon 2HB 96-well plates (Thermo Scientific, 3455) were coated with 50 ng/well of either *Py*CSP[NXA], *Py*CSP[ΔC], *Py*CSP[ΔN], or *Py*CSP[N2rC] constructs in 0.1 M NaHCO_3_, pH 9.5, overnight at room temperature. The plates were washed with PBS-containing 0.2% Tween-20 between each ELISA step. After coating, plates were blocked with 10% non-fat milk and 0.3% Tween-20 in PBS. RAM1 and 2F6 were serially diluted from 50 μg/mL to 5.12×10^−6^ μg/mL in PBS containing 10% non-fat milk and 0.03% Tween-20. Bound antibodies were detected with goat anti-mouse Ig-HRP (BD Biosciences, 554002) at 1:2000 dilution in PBS-containing 10% non-fat milk and 0.03% Tween-20. Plates were developed by adding 50 μL of TMB Peroxidase Substrate (SeraCare Life Sciences Inc, 5120-0083) to each well, incubating for 3 mins at room temperature, and stopping the reaction with 50 μL of 1 N H_2_SO_4_. Absorbance at 450 nm was measured by BioTek ELx800 microplate reader.

#### Repeat Peptides

RAM1 and 2F6 epitope mapping against repeat peptides was performed by Streptavidin capture ELISA. All incubations and washing steps were performed as described above. Immulon 2HB 96-well plates were coated with 50 ng/well of Streptavidin (NEB, #N7021S) in 0.1 M NaHCO_3_, pH 9.5, overnight at room temperature. Plates were blocked with 3% BSA in PBS, then coated with 250 ng/well of either biotinylated *Py*CSP[NXA] or biotinylated repeat peptides in 0.1 M NaHCO_3_, pH 9.5. A list of peptides used in this study and their sequences can be found in Table 1. Plates were then blocked again with 10% non-fat milk and 0.3% Tween-20 in PBS. After blocking, sRAM1 and 2F6 were serially diluted from 50 μg/mL to 5.12×10-7 μg/mL in PBS containing 0.2% BSA. Detection was performed with goat anti-mouse Ig-HRP as described above.

#### Mice

Female Swiss Webster (SW) mice (#032), 6-10 weeks of age, were purchased from Envigo. Female BALB/cJ mice (#000651), 6-8 weeks of age, were purchased from The Jackson Laboratory. Mice were housed under standard housing conditions and fed ad libitum water and food. Blood was collected by sub-mandibular, chin or cardiac puncture. All animal protocols were performed according to protocols reviewed by the Seattle Children’s Research Institute IACUC.

#### Generation of infected *Anopheles stephensi* mosquitoes

*P. yoelii* wild type strain 17×NL (BEI resources) was maintained in Swiss Webster mice (Envigo). Swiss Webster mice were injected intraperitoneal (IP) with 250 μL of infected blood at 3-5% parasitemia. Gametocyte exflagellation rate was checked 4 days post-injection. Mice were then anesthetized with 150 μL Ketamine Xylazine solution (12.5 mg/mL ketamine, 1.25 mg/mL xylazine in PBS) and naïve female *Anopheles stephensi* mosquitoes (3-7 days old) were allowed to feed on them. Mosquitoes were maintained at 23°C and 80% humidity with a 12L-12D light cycle. On day 10, the midguts from ten mosquitoes are dissected and oocysts counted as a measure of infection. On day 15 post-infection, mosquitoes are used for mosquito bite challenge or are dissected to isolate salivary glands and extract sporozoites.

#### *In vivo* protection studies

For passive transfer-challenge experiments, BALB/cJ mice were injected intraperitoneally 2F6 or RAM1, or 50C1, 24 h before mosquito bite challenge. Antibodies were diluted in PBS to final concentrations indicated in the Fig.s and injected into mice via a 25 5/8-gauge needle in volumes of 150 μg/ mouse. At 16-20 h following antibody injections, blood samples were taken via submandibular or chin bleeds (<50 μL/mouse) to assess antibody levels in the plasma prior to bite challenge. For experiments testing the effect of RAM1 on vaccination-induced protection, BALB/cJ mice were immunized with 20 μg of *Py*CSP[NXA] formulated with 20% v/v of Adjuplex (Invivogen, San Diego, CA, USA) as an adjuvant in PBS at 0, 2 and 6 weeks. Naïve mice served as controls. The vaccine formulation was administered intramuscularly to the hind leg in a volume of 50 μL/mouse. Six days after the third and final immunization, blood samples were collected via chin bleed to determine vaccine-induced anti-CSP titers and mice were then injected intraperitoneally with RAM1 or 50C1 at a dose of 500 μg/mouse. Mosquito bite challenge was performed 24 h following passive antibody transfer.

#### Mosquito Bite Challenge

Mice were anesthetized by IP injection of 150 μL Ketamine/Xylazine solution (12.5 mg/mL ketamine, 1.25 mg/mL xylazine in PBS). Once anesthetized, pairs of mice were each placed on a carton of 15 infectious mosquitoes and mosquitoes allowed to bite through the mesh top for 10 mins. Every thirty seconds mice were rotated among mosquito cartons to ensure equal exposure of mice and to maximize mosquito probing. Mice received subcutaneous PBS injections and recovered from anesthesia under a heat lamp. Mice were checked for the presence of blood stage parasites (patency) beginning 4 days post-challenge.

#### Assessment of blood stage patency

Patency was checked daily by blood smear beginning 4 days post-infection. Blood was collected by tail snip. Slides were fixed in methanol, dried, and then stained with giemsa (1:5 in H_2_O) for 10 mins. Slides were viewed at 100X and twenty fields of view examined for each smear. Mice were considered patent if 2 or more ring stage parasites were observed.

#### Sporozoite Quantification

To quantify the number of sporozoites per mosquito, we hand-dissected salivary glands from female *Anopheles* mosquitoes. These glands were then ground with a pestle and spun at 100x*g* to pellet debris. Sporozoites were counted on a hemocytometer.

#### Invasion assay

Freshly isolated sporozoites were exposed to 2F6 or RAM1 or 50C1 antibodies either alone or in combinations for 10 mins at indicated concentrations. 5×10^5^ Hepa1-6 cells were seeded in each well of a 24-well plate (Corning) and infected with antibody exposed *P. yoelii* sporozoites at a multiplicity of infection (MOI)=0.25 for 90 mins. After 90 mins of infection, cells were harvested with accutase (Life technologies) and fixed with Cytoperm/Cytofix (BD Biosciences). Cells were blocked with Perm/Wash (BD Biosciences) + 2% BSA for 1 h at room temperature then stained overnight at 4°C with antibodies to CSP- Alexafluor 488 conjugate. The cells were then washed and resuspended in PBS-containing 5 mM EDTA. Infection rate was measured by flow cytometry on an LSRII (Becton-Dickinson) and analyzed by FlowJo (Tree Star).

#### Traversal assay

Freshly isolated sporozoites were exposed to 2F6 or RAM1 or 50C1 antibodies either alone or in combinations for 10 mins at indicated concentrations. 5×10^5^ HFF-1 cells were seeded in each well of a 24-well plate (Corning). Cells were exposed to antibody treated sporozoites and high molecular mass Dextran-FITC (70 kDa) (Sigma) for 90 mins. Cells were harvested with accutase (Life technologies) and fixed with Cytoperm/Cytofix (BD Biosciences). Cells were blocked with Perm/Wash (BD Biosciences) supplemented with 2% BSA for 1 h at room temperature then stained overnight at 4°C with antibodies to CSP-Alexa fluor 488 conjugate. The cells were then washed and resuspended in PBS with 5 mM EDTA. Invasion rate and dextran uptake (traversal) were measured by flow cytometry on a LSRII (Becton-Dickinson) and analyzed by FlowJo (Tree Star).

#### Immunofluorescence

Freshly isolated sporozoites were exposed to 2F6 Alexa Fluor 405, RAM1- Alexa Fluor 594 or 50C1-Alexa Fluor 488 antibodies either alone or in pairwise combinations for 10 mins together or sequentially at indicated concentrations. Sporozoites were spun at 17000*g* for 5 mins and washed with 1xPBS-EDTA and fixed with 3.7% PFA for 20 mins. Sporozoites were transferred to 8-well chambered slides and the images were acquired with a 100×1.4 NA objective (Olympus) on a DeltaVision Elite High-Resolution Microscope (GE Healthcare Life Sciences). The sides of each pixel represent 64.5×64.5 nm and z-stacks were acquired at 300 nm intervals. Approximately 5-15 slices were acquired per image stack. For deconvolution, the 3D data sets were processed to remove noise and reassign blur by an iterative Classic Maximum Likelihood 95 Estimation widefield algorithm provided by Huygens Professional Software (Scientific Volume 96 Imaging BV, The Netherlands).

#### Statistical analysis

Statistical significance was determined with a one-way ANOVA for multiple comparisons or two-tailed unpaired student t-test. Analyses were performed using GraphPad Prism version 8 for Windows.

## Author contributions

K.V., R.C., O.T., S.L.B., N.D., M.Z., G.R.R.V., A.W., A.R., S.C., L.K.M., E.K.K.G., and R.P. performed the experiments. K.V., R.C., O.T., S.L.B., N.D, M.Z., G.R.R.V., A.K., and D.N.S analyzed the data. K.V., R.C., O.T., S.L.B., G.R.R.V., A.R., A.K., and D.N.S wrote the paper with input from all other authors. A.K. and D.N.S. supervised the research.

## Data and materials availability

The authors confirm that the data supporting the findings of this study are available within the article and its supplementary materials.

## Supplementary materials

**Supplementary Table 1:**
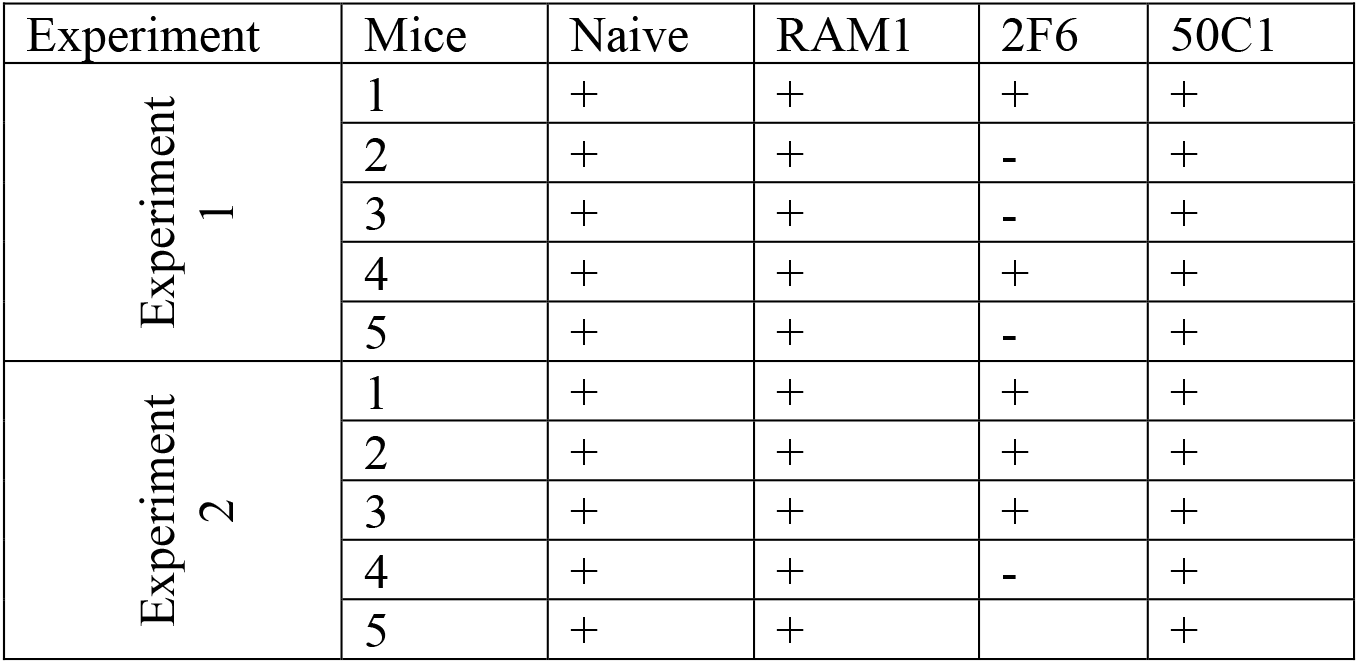
Patency status from two experiments following passive transfer of 2F6 or RAM1 or 50C1. + denotes infected; – denotes uninfected. Patency was assessed for 14 days post mosquito bite challenge. Related to Figure 4C.

| Experiment | Mice | Naive | RAM1 | 2F6 | 50C1 |
| --- | --- | --- | --- | --- | --- |
| Experiment 1 | 1 | + | + | + | + |
|  | 2 | + | + | - | + |
|  | 3 | + | + | - | + |
|  | 4 | + | + | + | + |
|  | 5 | + | + | - | + |
| Experiment 2 | 1 | + | + | + | + |
|  | 2 | + | + | + | + |
|  | 3 | + | + | + | + |
|  | 4 | + | + | - | + |
|  | 5 | + | + |  | + |

**Supplementary Table 2:**
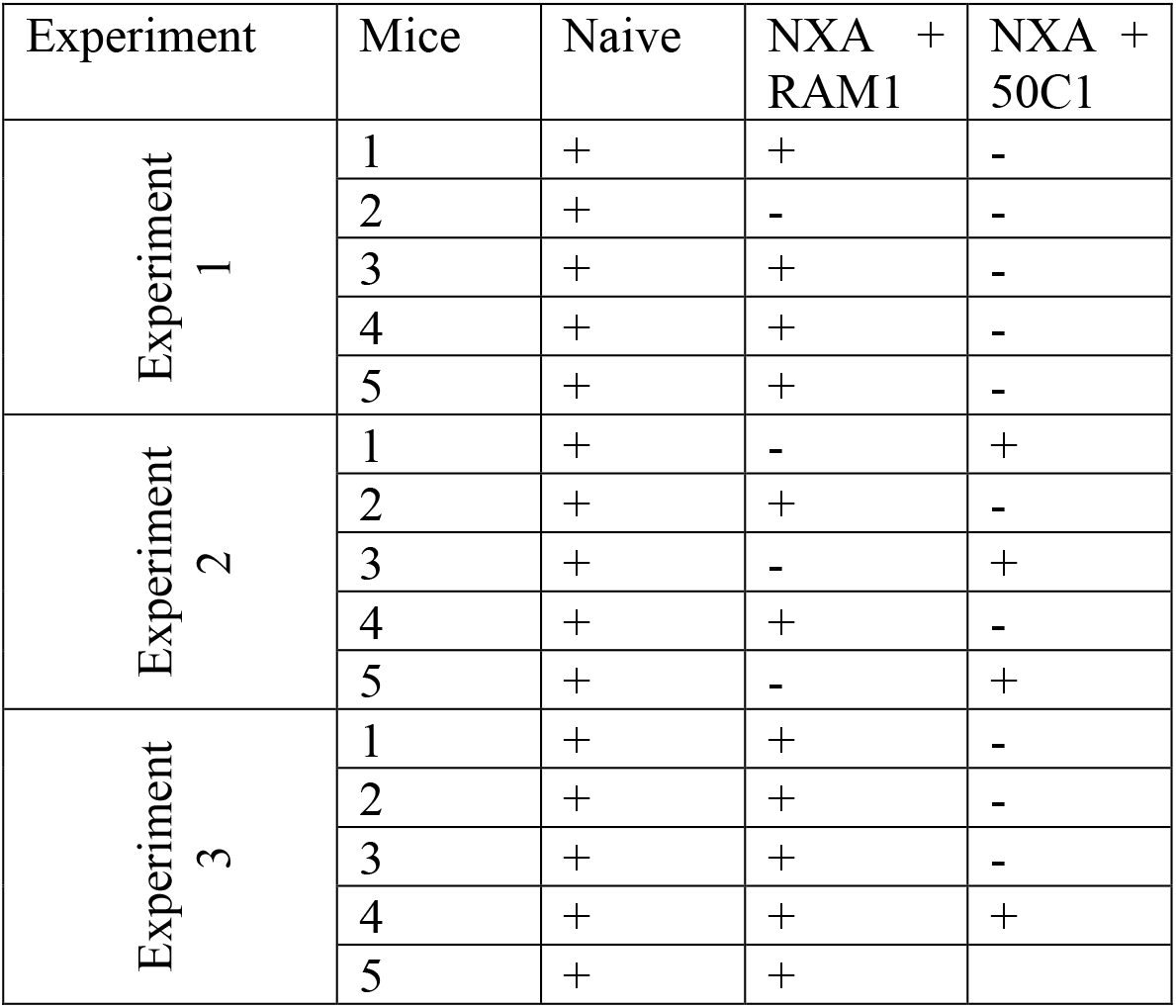
Patency status from all three experiments following active immunization with PyCSP[NXA] at 0, 2 and 6 weeks, followed by passive transfer of 2F6 or RAM1 or 50C1, a day prior to mosquito bite challenge. + denotes infected; – denotes uninfected. Patency was assessed for 14 days post mosquito bite challenge. Related to Figure 6D.

| Experiment | Mice | Naive | NXA +<br>RAM1 | NXA +<br>50C1 |
| --- | --- | --- | --- | --- |
| Experiment<br>1 | 1 | + | + | - |
|  | 2 | + | - | - |
|  | 3 | + | + | - |
|  | 4 | + | + | - |
|  | 5 | + | + | - |
| Experiment<br>2 | 1 | + | - | + |
|  | 2 | + | + | - |
|  | 3 | + | - | + |
|  | 4 | + | + | - |
|  | 5 | + | - | + |
| Experiment<br>3 | 1 | + | + | - |
|  | 2 | + | + | - |
|  | 3 | + | + | - |
|  | 4 | + | + | + |
|  | 5 | + | + |  |

